# Two spatially distinct posterior alpha sources fulfill different functional roles in attention

**DOI:** 10.1101/384065

**Authors:** R. Sokoliuk, S.D. Mayhew, K.M. Aquino, R. Wilson, M.J. Brookes, S.T. Francis, S. Hanslmayr, K.J. Mullinger

## Abstract

Directing attention helps to extract relevant information and suppress distracters. Alpha brain oscillations (8-12Hz) play a fundamental role in this process, with a power decrease facilitating processing of important information and power increase inhibiting brain regions processing irrelevant information. Evidence for this phenomenon arises from visual attention studies (Worden et al., 2000), however, the effect also exists in other modalities, including the somatosensory system (Haegens et al., 2011) and inter-sensory attention tasks (Foxe and Snyder, 2011). We investigated what happens when attention is divided between two modalities using both a multi- and unimodal attention paradigm while recording EEG over 128 scalp electrodes in two separate experiments. In Experiment 1 participants divided their attention between the visual and somatosensory modality to determine the temporal or spatial frequency of a target stimulus (vibrotactile stimulus or Gabor grating). In Experiment 2, participants divided attention between two visual hemifields to identify the orientation of a target Gabor grating. In both experiments, pre-stimulus alpha power in visual areas decreased linearly with increasing attention to visual stimuli. In contrast, alpha power in parietal areas showed lower pre-stimulus alpha power when attention was divided between modalities, compared to unimodal attention. These results suggest that there are two different alpha sources, where one reflects the ‘visual spotlight of attention’ and the other reflects attentional effort. To our knowledge, this is the first study to show that attention recruits two spatially distinct alpha sources in occipital and parietal brain regions, which act simultaneously but serve different functions in attention.

**SIGNIFICANCE STATEMENT:** Attention to one spatial location/sensory modality leads to power changes of alpha oscillations (^~^10Hz) with decreased power over regions processing relevant information and power increases to actively inhibit areas processing ‘to-be-ignored’ information. Here, we used detailed source modelling to investigate EEG data recorded during separate uni-modal (visual) and multi- (visual and somatosensory) attention tasks. Participants either focused their attention on one modality/spatial location or directed it to both. We show for the first time two distinct alpha sources are active simultaneously but play different roles. A sensory (visual) alpha source was linearly modulated by attention representing the ‘visual spotlight of attention’. In contrast, a parietal alpha source was modulated by attentional effort, showing lowest alpha power when attention was divided.

## INTRODUCTION

Allocation of attention helps extracting important and neglecting irrelevant information. Alpha brain oscillations (8-13Hz) potentially occupy this filtering role and lead to excitation or inhibition of sensory-specific regions, thereby facilitating or suppressing sensory processing (Klimesch et al., 2007; Jensen and Mazaheri, 2010; Mathewson et al., 2011). When attending to two spatial locations (right/left), a relative alpha power decrease is observed over brain regions processing relevant information compared with regions inhibiting irrelevant information. Such a hemispheric alpha power lateralization over occipito-parietal regions has been shown many times in visuospatial attention (Foxe et al., 1998; Worden et al., 2000; Zumer et al., 2014). This has also been observed in the somatosensory system (Anderson and Ding, 2011; Haegens et al., 2011, 2012; van Ede et al., 2011) and in inter-sensory attention (Foxe and Snyder, 2011; Gomez-Ramirez et al., 2011; Bauer et al., 2012).

What happens if attention is divided between two sensory modalities simultaneously? Would this provoke an alpha-power-imbalance between sensory-specific regions reflecting the peak location of attention, like recently observed for spatially divided visual attention (Gould et al., 2011)? Existing literature showed evidence for alpha-power-modulation over sensory-specific brain regions, however, attention was not divided between two senses simultaneously (Foxe and Snyder, 2011; Gomez-Ramirez et al., 2011; Bauer et al., 2012). Functional magnetic resonance imaging (fMRI) evidence suggests attention also modulates activity over higher-level frontal and parietal areas (Corbetta and Shulman, 2002) that modulate lower-level sensory regions via top-down-control (Bressler et al., 2008). Inhibiting frontal eye field (FEF) and inferior parietal lobule (IPL) using repetitive Transcranial Magnetic Stimulation (rTMS), Capotosto et al. observed increased reaction times and decreased accuracy for visual detection and thereby confirmed fMRI results. They concluded that inhibiting these regions disrupted the control over visual alpha oscillations and altered behaviour (Capotosto et al., 2009). According to the authors, both primary sensory and parietal regions are important for controlling attention allocation. Hints of this in EEG are shown by the spatial and functional dissociation of occipital and parietal alpha sources during visual perception (Gulbinaite et al., 2017).

Here, we investigated i) potential differences in the role of alpha oscillations between uni- and multi-modal attention and ii) the effect of dividing attention between two sensory modalities. We used multi-modal (Experiment 1) and uni-modal (Experiment 2) attention paradigms while recording scalp EEG over 128 electrodes.

In Experiment 1 (Figure 1 A), participants divided their attention between the visual and somatosensory system and judged whether visual/vibrotactile stimuli were targets (low or high spatial/temporal frequency) or distracters (medium spatial/temporal frequency) and reported the target frequency. Attention was manipulated using a pre-stimulus visual cue with varying levels of certainty (see Figure 1) before a target was presented in one modality simultaneously to a distracter in the other. Experiment 2 (Figure 1 B) manipulated attention between visual hemifields (cf. Posner et al., 1980) using a pre-stimulus visual cue with varying levels of certainty (see Figure 1; Gould et al., 2011). Participants identified and reported the orientation of visual Gabor gratings. Targets (horizontal/vertical gratings) and distractors (diagonal gratings) were presented simultaneously to each side of a central fixation cross.

**Figure 1:**
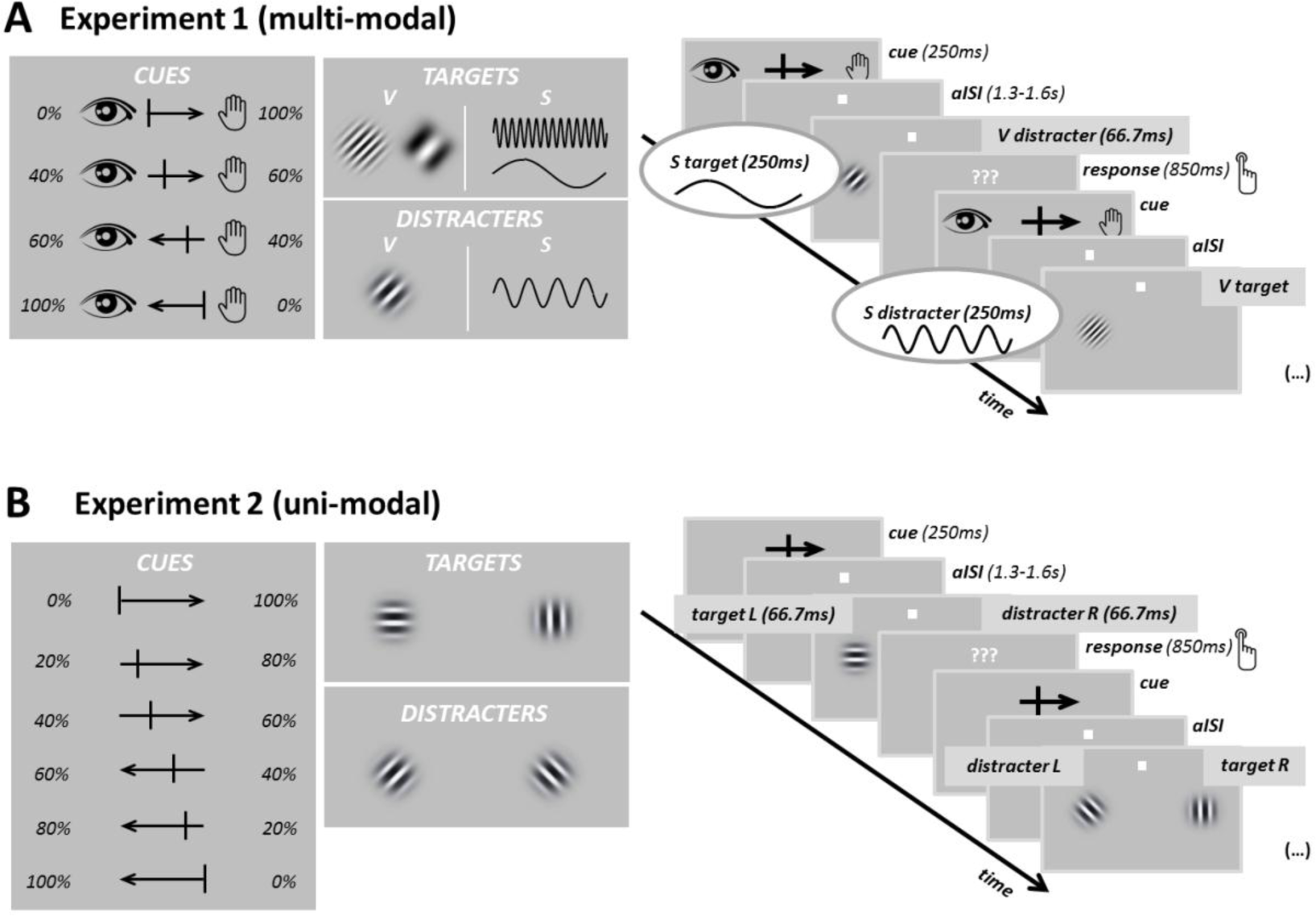
Paradigms of Experiment 1 and Experiment 2. (**A**) shows the attention paradigm used in Experiment 1. The left panel shows the attentional cues used to manipulate participants’ attention for the four different attention conditions. The eye represents “attention to the visual system” while the hand represents “attention to the somatosensory system”. The arrows point in the direction of the modality that should be more strongly attended to. The numbers (e.g. 0%) were not presented during the experiment but are shown here for clarity. Target stimuli in the visual domain were high and low frequency Gabor patterns whereas stimuli with a medium spatial frequency represented visual distracters (see middle panel). In the somatosensory domain, stimuli showing a high or a low temporal frequency served as targets whereas medium frequency stimuli were distracters (see middle panel). On the right side, the temporal sequence of the experiment is shown. A cue was presented for 250ms before a blank screen only showing the fixation point for 1.3-1.6s (aISI). Then both, visual (66.7ms) and somatosensory stimuli (250ms) were presented simultaneously, while only one of them represented the target stimulus. Subjects then had 850ms to respond whether the target was high or low frequency before the next trial. (**B**) The left panel shows the visual cues used to manipulate participants’ attention in the six attention conditions of Experiment 2. The arrows are pointing towards the side of the visual field to which more attention should be paid with dividing lines indicating how attention should be divided (as in Experiment 1). Again numbers (e.g. 0%) are only shown for clarity and were not presented. As in Experiment 1, each trial started with the presentation of a visual cue (250ms) before a blank screen with only the fixation point was presented for 1.3 – 1.6s (aISI), see right panel. Then, stimuli appeared on both sides of the visual field whereat only one of them was a target whose orientation (e.g. “horizontal or vertical”, see middle panel) had to be reported within 850ms before the next trial started. The middle panel showing target and distractors is an example which was used for half the subjects; for the other half the subjects the target and distractors were the opposite. Note: to facilitate visibility in these schematics, the visual stimuli are larger than the actual size these stimuli occupied on the screen in the experiment.

A Linearly Constrained Minimum Variance (LCMV) beamformer (Van Drongelen et al., 1996) source localised changes in pre-stimulus alpha power. Two alpha sources were identified in Experiment 1: A visual source decreased linearly in power with increasing attention to visual stimuli; a second source in the parietal cortex modulated by task difficulty showed lower alpha power when attention was divided between modalities. Experiment 2 shared the visual source with linear attention modulation however parietal brain regions were not as strongly modulated.

To our knowledge, this study is the first to reveal two spatially distinct alpha mechanisms acting simultaneously and yet performing different roles in attention: a sensory, visual alpha source reflecting the current location of attention and a parietal alpha source modulated by task difficulty and reflecting attentional effort.

## METHODS

### Participants

Data were acquired from 20 healthy participants (all right-handed, 10 females, mean age 28.1 ± 3.8 years) with normal or corrected to normal vision. One participant was not included in final data analysis because of the absence of an anatomical MRI scan that prohibited complete data analysis. 15 out of these 20 participants performed two attention paradigms (Experiment 1 and Experiment 2), the remaining four subjects only participated in Experiment 1.

The study was approved by the University of Birmingham Research Ethics Committee. Before the start of the experiment(s), participants provided informed written consent.

### Stimuli and Task

Visual and somatosensory stimuli were presented using Psychophysics Toolbox (Version 3; Brainard, 1997) running in MATLAB (version 2014b; MathWorks) on a desktop computer (Windows 7). Participants sat comfortably in a dark room. To minimize head movement and maintain a constant degree of visual angle for the visual stimuli, their head was kept stable using a chin rest.

Visual stimuli were presented in Experiments 1 and 2 on a grey background. Gabor gratings were presented briefly (presentation time: 66.7ms; radius: 1.75 degrees of visual angle, phase: 180°), on a grey background at a distance of 57cm, using a cathode ray monitor (resolution: 600 x 800 pixels). These stimuli were presented vertically centered and with a horizontal eccentricity of ±8 degrees of visual angle from a horizontally centered white fixation point (radius: 0.1 degrees of visual angle). In Experiment 2, two Gabor gratings were presented to the left and right of the fixation point whereas in Experiment 1 a single visual stimulus was presented to the left of the fixation point, simultaneously with a 250ms long vibrotactile stimulus to the tip of the left index finger using a piezoelectric stimulator (Dancer Design, St. Helens, United Kingdom, http://www.dancerdesign.co.uk).

In Experiment 1 the multimodal attention task was conducted (see Figure 1A). While fixating on the fixation cross, subjects had to covertly divide their attention between two sensory modalities, attending either more to visual or more to somatosensory stimuli (0/100% or 40/60% attention towards somatosensory/visual domain and vice-versa). A visual cue (5×2.5 degrees of visual angle) was presented at fixation at the beginning of every trial for 250ms, indicating how attention was to be divided. Cues took the form of black arrows indicating the likelihood of subsequent target appearance in each modality (cf. Figure 1A). After an asynchronous inter-stimulus interval (aISI) of 1.3-1.6s (aISIs were randomly chosen for every trial reaching from 1.3s (minimum aISI) to 1.6s (maximum aISI)), during which participants were required to divide their attention between modalities according to the pre-stimulus cue, visual and somatosensory stimuli were presented simultaneously. Gabor patterns were presented in a tilted orientation: for half of the participants they were tilted at 45°, for the other half at −45°. Stimuli with a low or high spatial frequency (0.025 cycles/pixel and 0.1 cycles/pixel) were visual targets and medium frequency stimuli (0.05 cycles/pixel) were visual distracters. In the somatosensory domain, vibrotactile stimulation at a low or high temporal frequency (4 Hz and 52 Hz) served as somatosensory targets and those at medium temporal frequency (16 Hz) as somatosensory distracters. In every trial, one target (e.g. a visual Gabor pattern with a high spatial frequency) and one distracter (e.g. a somatosensory stimulus with a medium temporal frequency) stimulus were presented simultaneously. After stimulus presentation, white question marks (5×1.5 degrees of visual angle) indicated an 850ms response period where participants pressed a button with their right index finger to report the frequency of the target (two different keys: high or low frequency, regardless of probed modality) as quickly as possible.

In Experiment 2 the uni-modal attention task was conducted (see Figure 1B). This second experiment had a similar structure to Experiment 1 but used only visual Gabor gratings (spatial frequency: 0.05 cycles/pixel), akin to a classic Posner task (Posner et al., 1980). Subjects had to covertly direct their attention in a graded fashion either more to the left or more to the right visual hemifield (0/100%, 20/80%, or 40 /60%, attention towards left/right visual hemifields and vice-versa) while they fixated on a central fixation cross (similar to Gould et al., 2011). Trials started with the presentation of a visual cue (5×2.5 degrees of visual angle; presentation time: 250ms) in the form of black arrows indicating where subjects should direct their spatial attention (cf. Figure 1B). As in Experiment 1, this was followed by an aISI of 1.3-1.6s, before visual stimuli were presented to the left and right of the fixation point. For half of the participants, horizontal and vertical gratings were target stimuli and rightwards (45°) and leftwards (−45°) tilted gratings served as distracters, while for the other half of participants the opposite was true. In every trial one target (e.g. horizontal grating) and one distractor (e.g. rightwards tilted grating) appeared simultaneously at opposite sides of the fixation cross. After stimulus presentation, a white question mark was presented for 850ms to indicate the response period. The task was to respond as fast as possible to indicate the orientation of the target grating (two different keys: e.g. horizontal or vertical).

In both experiments, participants were given feedback on their performance (accuracy and reaction time) which was displayed after each experimental run to maintain their motivation for performing the tasks.

Participants completed a training run consisting of 10 trials per attention condition (resulting in a total of 40/60 trials for Experiments 1/2, respectively) before they performed the same task in a staircase experimental run, where the contrast of the visual stimuli was adapted according to participants’ performance (60 trials per attention condition) to ensure an accuracy of ^~^80% was achieved. For somatosensory stimuli, a similar procedure was used to adapt the amplitude of vibrotactile pulses.

The subjects then started the experiment and performed 150 trials per attention condition giving a total of 600/900 trials for Experiments 1 and 2, respectively. Experiments were divided into 3 individual runs; all runs contained equal number of trials of each attention condition (50 trials/condition/run). All trials of a given attention condition within a run were grouped together in one block, the order of the blocks between runs was varied pseudo-randomly. The whole study took ^~^1.5 hours per participant, including short breaks that the participants took between runs.

### EEG data acquisition

EEG data was recorded from 128 active scalp electrodes following an equi-radial montage at 1024 Hz sampling rate using a Biosemi EEG system (Amsterdam, Netherlands) with a reference electrode (common mode sense electrode) placed parieto-centrally for the recording. In addition, EOG was recorded using 3 active ocular electrodes with the horizontal electrodes being placed near the two temples and the vertical electrode below the left eye. In Experiment 1, data was recorded in 3 runs of ^~^9 minutes each, in Experiment 2 the three runs consisted of ^~^12 minutes each.

After each EEG recording session, the individual electrode positions were digitised relative to the surface of the head with a Polhemus FASTRAK using Brainstorm software (Tadel et al., 2011) running in Matlab (MathWorks). In addition, each subject attended a separate MRI session where a T1-weighted anatomical image (MPRAGE sequence) of the head, including the nose, with 1mm isotropic resolution was acquired on either a 3T or 7T MRI system which was registered with the digitised head shape.

### Data analysis

#### Behavioural

Behavioural parameters analysed were reaction time and accuracy. In order to analyse significant differences between attention conditions, a repeated measures 2-way ANOVA was computed for both behavioural parameters and experiments separately, with factors: (i) attention condition (60 and 100% for Experiment 1 and 60, 80, and 100% for Experiment 2), and (ii) attended modality (somatosensory and visual) or hemifield (left and right). Post-hoc paired sample t-tests were used to identify individual differences between attention conditions, and p-values were subsequently Bonferroni-corrected to account for multiple comparisons.

#### EEG

All EEG data processing was carried out using the Matlab toolbox Fieldtrip (Oostenveld et al., 2011).

##### Pre-processing

Data were read in as continuous data, for each channel data were notch filtered (49-51Hz) to reduce line noise, detrended to remove linear drifts and demeaned (subtracting the average signal recorded over the whole time course at each channel) to remove between run baseline effects. By visual inspection, noisy channels (i.e. channels with obvious artifacts) were removed from further data analysis. This resulted in a group mean of (±standard error (SE)) 117 ± 4 / 116 ± 5 channels remaining for further analysis for Experiment 1/2 respectively. Independent component analysis (ICA, logistic infomax ICA algorithm, (cf. Bell and Sejnowski, 1995) was then performed to discard eye blinks from the recorded data, with an average of 1 ± 0.6 ICs for Experiment 1 and 1.5 ± 1.5 ICs for Experiment 2 removed from each data set. The remaining ICs were re-projected to the channel level. Finally, data were re-referenced to the average of all the non-noisy channels that remained for each subject and run.

These data were subsequently used for time-frequency analysis on the sensor and source level.

##### Sensor level analysis

Data were epoched into 1.7s (−1.5s until +0.2s relative to the stimulus presentation onset) segments for every trial and the separate runs of the experiment concatenated. All trial level data were visually inspected and noisy trials removed for each subject, resulting in 818±12.4 / 539±11.7 (number of trials ± standard error of the mean [SEM]) trials of data remaining for Experiments 1/2. Furthermore, those trials where the subject had responded incorrectly to the target were subsequently removed such that 727±16.5 / 471±15.6 trials remained for Experiment 1/2. Further analysis details and results from the sensor level are shown in the Supplementary Material, as the focus of this work was on the source analysis.

##### Source level analysis

Individual, 4-layer (scalp, skull, CSF, & brain) boundary element (BEM) head models were constructed from the individual subject T_1_-weighted anatomical images using the Fieldtrip toolbox with the ‘dipoli’ method (http://www.ru.nl/neuroimaging/fieldtrip) (Oostenveld et al., 2011). Individual electrode positions were aligned to the scalp surface of the subject’s T_1_ using the fiducial points and headshape to inform alignment. In 4 of the 19 participants, no individual electrode positions were recorded due to technical problems; therefore, in these subjects the average electrode positions of the 11 other participants sharing the same electrode layout were used and warped to the scalp surface extracted from the segmented individual T_1_-weighted scans.

Beamforming analysis was performed using a Linearly Constrained Minimum Variance (LCMV) beamformer (Van Drongelen et al., 1996; Van Veen et al., 1997; Robinson and Vrba, 1999) implemented in the Fieldtrip toolbox, to spatially localize changes in alpha power between different attention conditions. The continuous data for each run were first filtered into the alpha frequency band (8-13Hz) and then epoched −1.5s to +0.2s relative to stimulus onset. The noisy and incorrect response trials, identified from the broadband visual data inspection (see “Sensor level” section above) were removed. Remaining trials were then concatenated over runs, downsampled to 500 Hz and beamformer weights (also known as a spatial filter) (Van Veen et al., 1997) derived. All attention conditions within an experiment were considered together to calculate these weights as the spatial sources of the alpha power were not hypothesized to change between conditions but only their relative amplitude.

For each subject the preprocessed, cleaned and downsampled sensor level data were then separated into trials for each of the attention conditions. The number of trials in each condition was reduced to match that of the condition with the minimum number of trials remaining. This data rejection process was done by randomly removing trials from conditions containing more trials than the minimum. This process ensured all source localization comparisons were performed on equal amounts of data to avoid biases. An average of 105± 22 of the 150 trials per condition for Experiment 1 and 107 ± 16 of the 150 trials per conditions for Experiment 2 remained (mean ± SE over subjects) for further source analysis.

To enable alpha power to be calculated only during the aISI, trials were then segmented resulting in a time window from −1.3s to 0s relative to stimulus onset and concatenated together for each condition to ensure no baseline effects within trials were removed. The source power at each location in the brain BEM (0.5 cm grid) was estimated for each condition, using the previously derived weights from all conditions. These source power maps were then used to calculate the alpha modulation index (AMI) source maps for both experiments for each subject using Equation 1, where the source power estimates at each location in the brain for each condition were input, as previously employed (Zumer et al., 2014).

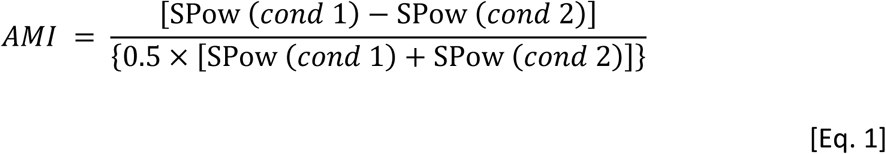

In Experiment 1, the AMI between trials where participants focused on one modality compared to focusing on the other, e.g. between 100% attention to the visual domain vs. 100% to the somatosensory domain, was calculated using Equation 1, where SPow (‘**S**ource **P**ower’) was calculated for every location in the brain (on the 0.5 cm grid) and is the power estimate of the alpha band signal over the time period −1.3 to 0s relative to stimulus onset for all trials in a given condition. Here, *cond 1* denotes attend 100% to visual (and 0% to somatosensory) stimuli whilst *cond 2* denotes attend 100% to somatosensory (and 0% to visual) stimuli.

Furthermore, the AMI between trials where participants focused on one modality (100% visual or somatosensory; *cond 1* in Equation 1) and those where attention was divided between modalities (60% visual (i.e. 60% visual and 40% somatosensory) or somatosensory (i.e. 60% somatosensory and 40% visual); *cond 2* in Equation 1) was computed. The equivalent AMIs were calculated for Experiment 2. First, attention conditions 100% left (*cond 1* in Equation 1) and 100% right (*cond 2* in Equation 1) were compared. Then trials were compared according to whether subjects paid attention to only one side of the visual field (100%; *cond 1* in Equation 1) or divided their attention between left and right hemifields (60%; *cond 2* in Equation 1).

The AMI(100%,100%) contrasts “*100% visual (V) vs. 100% somatosensory (S)*” and “*100% left (L)* vs. *100% right (R)*” for Experiments 1 and 2 respectively, were designed to investigate differences in alpha modulation depending on the attentional cue. Whilst the AMI (100%,60%) contrasts “*100% (visual/somatosensory)* vs. *60% (visual/somatosensory)*” and “*100% (left/right)* vs. *60% (left/right)*” for Experiments 1 and 2 respectively, were designed to elucidate whether task difficulty was reflected by modulations in alpha power.

AMI source maps for each subject were spatially normalized to the MNI template before being averaged over subjects for each experiment to provide a grand average. The different grand average AMI source maps were visually inspected for local minima and maxima for the two experiments. In both experiments, local minima and maxima were observed over the visual cortex (AMI(100%,100%)) and the parietal cortex (AMI(100%,60%)), respectively. For Experiment 1, all stimuli were presented on the left thus hypothesized to recruit the right hemisphere of the brain primarily. Therefore, the maximum AMI value peak location in the right parietal cortex (anatomically defined) from the AMI(100%,60%) maps and a minimum AMI value peak location in the right visual cortex from the AMI(100%,100%) were found for each subject individually. For Experiment 2 bilateral stimulus presentation resulted in hypothesized responses in both hemispheres. Therefore the AMI maxima were identified in the right and left parietal cortices (AMI(100%,60%)), and in the left visual cortex (AMI(100%,100%)). Furthermore, the AMI minimum in the right visual cortex was identified (AMI(100%,100%)). All peak locations within the anatomically defined regions were identified for each subject individually.

##### Peak location analysis

The identified peak locations were used as virtual electrode (VE) locations from which alpha frequency time courses were extracted for each participant individually. Time courses were obtained at each VE location by multiplying the cleaned, continuous, downsampled channel level data (used to derive the initial weights) by the respective alpha beamformer weights derived over all data (see above). Time courses were then demeaned before a Hilbert transform was performed to provide a measure of alpha power at each VE location interrogated for each subject. The data were then epoched −1.3s to 0s relative to stimulus onset (i.e. the aISI period) and separated into conditions (using the same balancing procedure used to derive the source maps). The average alpha power over trials for each condition was found and then averaged over the aISI period (−1.3 – 0s) to provide a measure of mean alpha power per condition in the visual and parietal cortices.

For Experiment 2, data from left and right hemispheres were combined by flipping the attention conditions (attention left 100% = attention right 100% etc.) for the data recorded over the right hemisphere, effectively resulting in alpha power modulations from the left parietal and visual cortices (cf. Waldhauser et al., 2016). This procedure was designed to increase signal to noise.

To take account of between subject variance, alpha power values were then normalised by the maximum average alpha power value in any condition for each subject. Subsequently, the grand average over subjects was computed. These were tested for linear and quadratic modulation over conditions by fitting the data first to linear and then to quadratic functions using the Matlab function *polyfitn*.

##### Automated anatomical labeling (AAL) analysis

To test whether the linear and quadratic modulations observed from the peak location analysis were statistically significant, we performed additional analyses based purely on anatomically parcellated brain regions and therefore not biased by the AMI source maps in identification of locations to interrogate. Brain regions were parcellated using the automated anatomical labelling (AAL) atlas (Tzourio-Mazoyer et al., 2002). Only the anatomical regions in which an alpha related response was predicted were interrogated. Therefore 15 AAL regions in the right hemisphere, spanning from the visual cortex to the somatosensory cortex and 26 AAL regions in right and left hemisphere, reaching from visual to parietal cortex, were investigated in Experiments 1 and 2 respectively (see table 1 and 2).

**Table 1:**
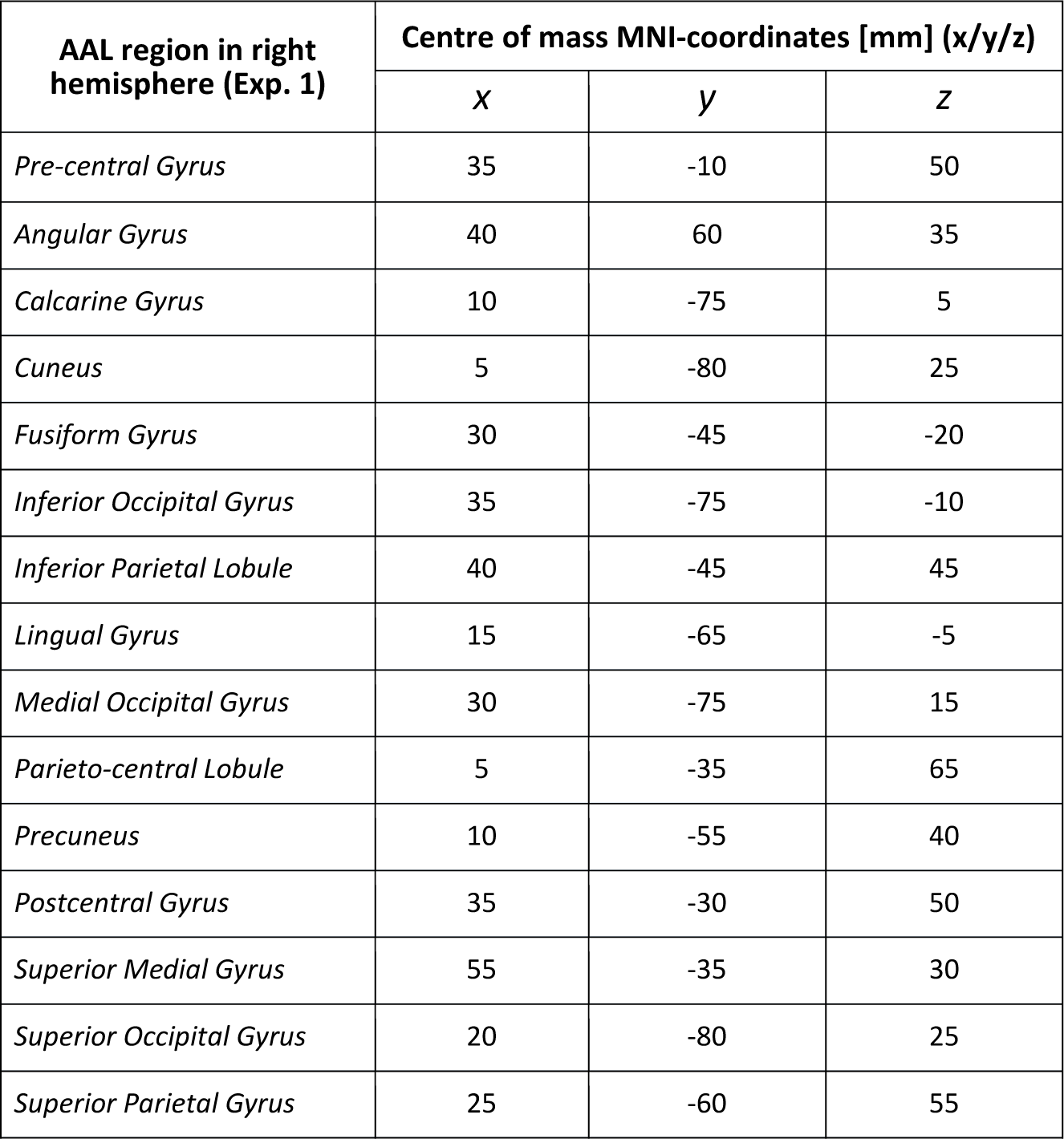
AAL regions with MNI coordinates of centre of mass investigated in Experiment 1.

**Table 2:** AAL regions with MNI coordinates of centre of mass investigated in Experiment 2.

| AAL region in right and left hemisphere (Exp.2) | Centre of mass MNI-coordinates [mm] (x/y/z) |  |  |
| --- | --- | --- | --- |
|  | x | y | z |
| *R/L Angular Gyrus* | 40/-40 | 60/-60 | 35/35 |
| *R/L Calcarine Gyrus* | 10/-15 | -75/-75 | 5/10 |
| *R/L Cuneus* | 5/-15 | -80/-80 | 25/25 |
| *R/L Fusiform Gyrus* | 30/-35 | -45/-45 | -20/-20 |
| *R/L Inferior Occipital Gyrus* | 35/-35 | -75/-80 | -10/-10 |
| *R/L Inferior Parietal Lobule* | 40/-40 | -45/-45 | 45/50 |
| *R/L Lingual Gyrus* | 15/-15 | -65/-65 | -5/-5 |
| *R/L Medial Occipital Gyrus* | 30/-35 | -75/-75 | 15/20 |
| *R/L Postcentral Lobule* | 10/-5 | -25/-35 | 65/65 |
| *R/L Precuneus* | 10/-10 | -55/-55 | 40/40 |
| *R/L Superior Medial Gyrus* | 55/-50 | -35/-35 | 30/35 |
| *R/L Superior Occipital Gyrus* | 20/-25 | -80/-75 | 25/30 |
| *R/L Superior Parietal Gyrus* | 25/-25 | -60/-55 | 55/55 |

The following analysis approach was used, as has been previously employed on MEG data (Brookes et al., 2016). For each subject, all AAL regions were warped onto the individual subject’s T_1_-weighted image and timecourses were then extracted from all VE locations (on a 0.5cm grid) which fell within the AAL regions. The VE time courses were extracted using the same data and processes used for the *peak location analysis*. Time courses from VE locations (each grid point) were weighted according to the Euclidian distance of the VE location to the centre of gravity of the respective AAL region. After applying the correct weighting, time course data were summed over all VEs per AAL region, to give one time course per AAL region containing all trials, which was then demeaned. The Hilbert transform was subsequently applied to time courses for each AAL region. The data were then epoched-1.3s to 0s relative to stimulus onset (i.e. the aISI period) and separated into conditions (using the same trial balance used for the source maps and peak responses). The alpha power time courses for each AAL region were then averaged over trials and aISI time window within each attention condition and subject. The outcome of this processing was 15×4 (Experiment 1: 15 AAL regions and 4 attention conditions) or 26×6 (Experiment 2: 26 AAL regions – including AAL regions in the left hemisphere but excluding AAL regions within the somatosensory cortex – and 6 attention conditions) alpha power values per subject. Data of Experiment 2 was averaged between AAL regions across hemispheres by flipping the attention conditions, resulting in 13 AAL datasets per subject.

Before averaging over subjects, the resulting 4/6 alpha power values for the attention conditions in Experiment 1/2 per AAL region were normalized by the alpha power value of the attention condition that showed the maximum power, removing between subject variance to ensure between condition variance was interrogated. Given the apparent linear and quadratic modulation patterns derived from the peak location analysis, the normalised average alpha power over all subjects (i.e. 15/19 data points per condition for Experiments 1/2, respectively) were first fit with a linear function and subsequently, if no linear modulation was observed, with a quadratic function across the attention conditions. Significance of the fits obtained on the real data was determined through Monte Carlo permutation tests (25,000 repetitions). Here, for every AAL region, the real data fits were compared with surrogate distributions of linear and quadratic terms of the respective AAL regions, derived from shuffling data between the different attention conditions for every subject individually and performing new linear and quadratic fits over the 4/6 surrogate attention conditions. The p-values obtained were then corrected for multiple comparisons (i.e. AAL regions) using False Discovery Rate (FDR) correction (Benjamini and Hochberg, 1995; Yekutieli and Benjamini, 1999).

## RESULTS

### Behaviour

#### Experiment 1 (multimodal task)

A 2-way repeated measures ANOVA with main factors of cue (100 or 60% attention) and modality (attention to visual or somatosensory modality) revealed that the accuracy for detecting the spatial/temporal frequency of target Gabor gratings/vibrotactile stimuli was significantly higher in the “attend 100%” condition than in the “attend 60%” condition (p-value = 1.3×10^−7^; F = 34.3, Figure 2A, upper panel). No significant effect of modality (p-value = 0.21; F = 1.6) and no interaction between cue and modality was observed (p-value = 0.4; F = 0.67; Figure 2A, upper panel).

**Figure 2:**
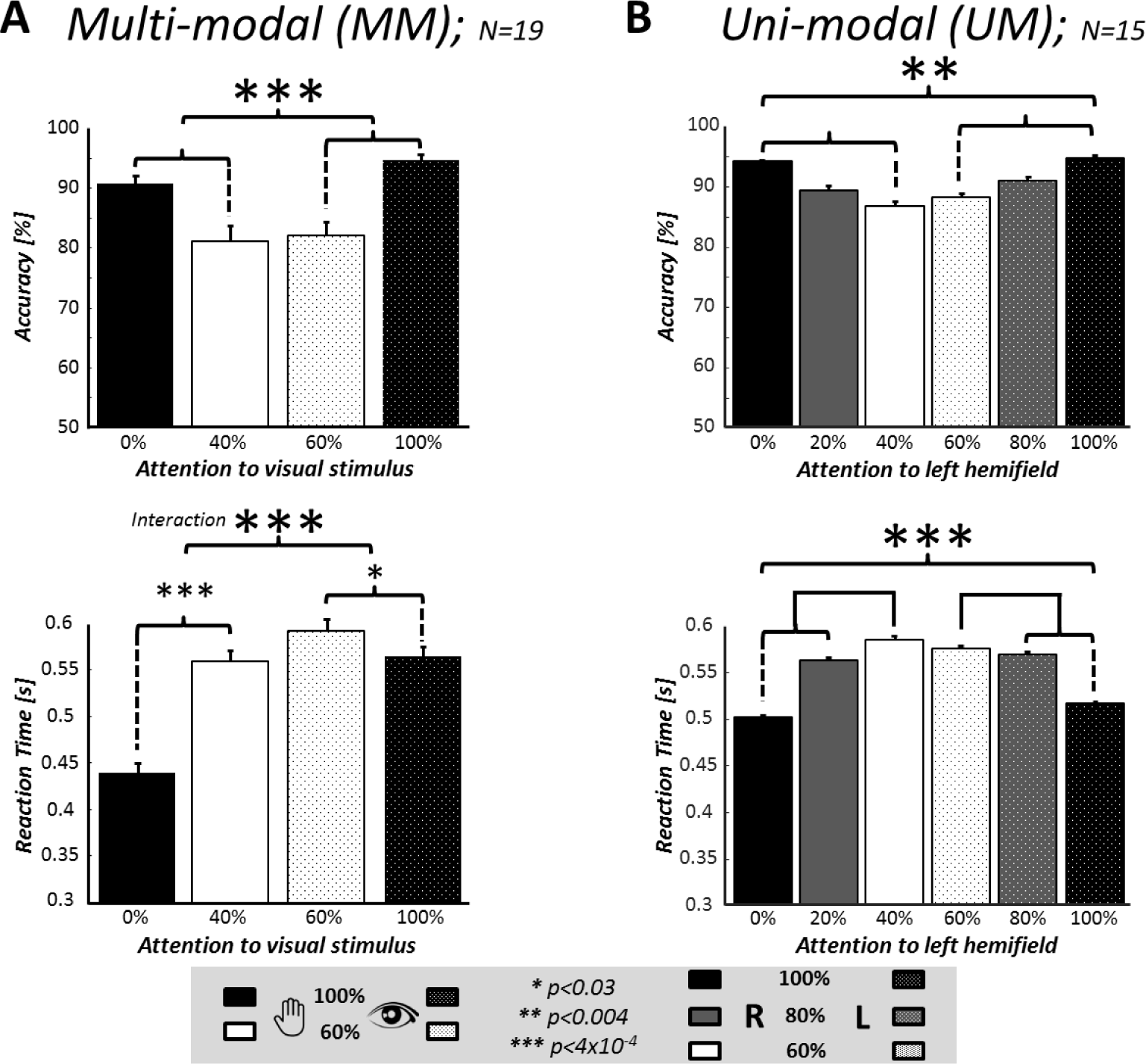
Behavioural measures of accuracy (top panels) and reaction time (bottom panels) across attention conditions. Panel A shows the behavioural results of the multimodal (visual vs somatosensory) paradigm (upper panel: accuracy achieved in each condition, lower panel: reaction times). A significant interaction between cue and attended modality in the reaction time shows that participants’ behaviour is modulated to a greater extent when attention is directed to the somatosensory modality (0 and 40%) than the visual modality (60 and 100%). Panel B shows behavioural data for the uni-modal (visual) paradigm (upper panel: accuracy achieved; lower panel: reaction times). All bars denote the mean response over subjects whilst error bars denote the SEM over subjects. Asterisks denote p-values from 2-way ANOVAs (see legend).

A second 2-way repeated measures ANOVA compared the reaction times (RTs) across attention conditions. With main factors of cue and modality, this test revealed a main effect of cue (p-value = 1.1×10^−8^; F=41.8). A significant interaction between factors cue and modality (p-value = 1.03×10^−4^; F = 16.9) revealed a stronger effect of cue on RTs when subjects attended to the somatosensory stimuli (Figure 2A, lower panel).

#### Experiment 2 (unimodal task)

A 2-way repeated measures ANOVA with main factors of cue (60, 80, and 100% attention) and side of presentation (left or right visual hemifield) revealed as the main effect that accuracy for detecting the orientation of target Gabor gratings was significantly higher in the “attend 100%” than in the “attend 60%” condition (p-value = 3.37×10^−4^; F = 8.8; Figure 2B, upper panel). No significant effect of side of presentation (p-value = 0.63; F = 0.2) and no interaction between cue and side of presentation was observed (p-value = 0.64; F = 0.4; Figure 2B, upper panel).

A further 2-way repeated measures ANOVA with main factors of cue and side of presentation showed that RT was significantly shorter when subjects only attended to one side of the visual field (100 vs. 0% attention), than when they divided their attention between hemifields (80 vs 20% and 60 vs. 40% attention; p-value = 4.1×10^−6^; F = 14.4). There was no significant interaction between cue and side of presentation (p-value = 0.3; F = 1.1; Figure 2B, lower panel).

### EEG responses

As there was hypothesized to be more than one alpha power response from different cortical areas, we focus the results presented on the source level results where spatial localisation aids interpretation of the data.

#### Experiment 1

In order to investigate potential differences in alpha power between the attention conditions, we first compared trials where subjects only paid attention to visual stimuli (100% V; *cond 1* in Equation 1) with trials where they only attended to somatosensory stimuli (100% S; *cond 2* in Equation 1). We computed the alpha modulation index (‘AMI’; Equation 1) on the beamformer results which revealed a negative response in right visual cortex, indicating an alpha power decrease in visual cortex with increasing attention to the visual domain as shown in Figure 3A. No alpha power modulation was found in somatosensory areas between these two attention conditions (Fig 3A). The AAL analyses supported this observation revealing a significant linear modulation of alpha power (p-value = 0.02, fdr-corrected) observed in the right inferior occipital gyrus, see Figure 3B.

**Figure 3:**
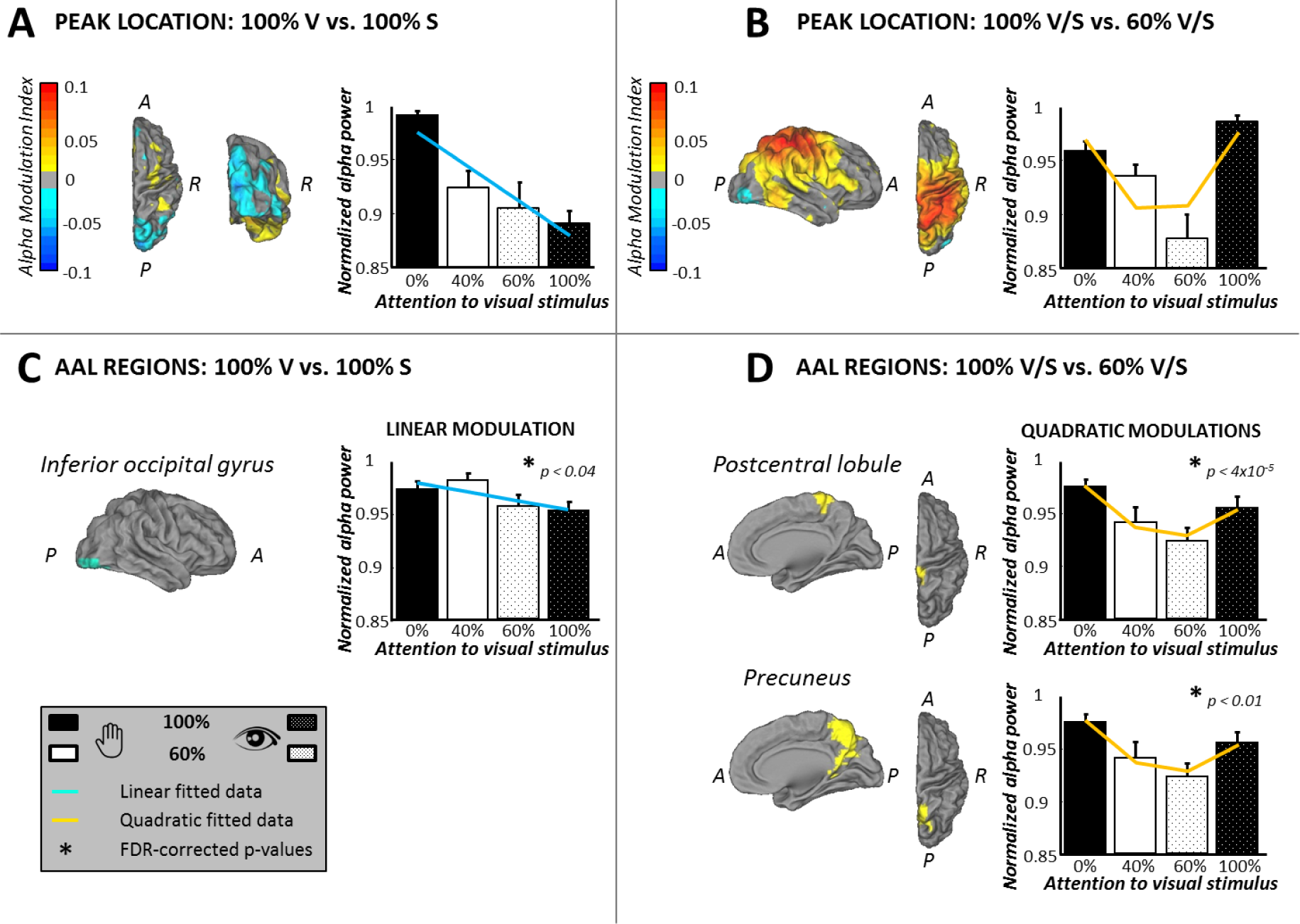
Source analysis results of Experiment 1. **Panel A** shows source analysis results for Experiment 1 when contrasting the conditions 100% visual (0% somatosensory) with 100% somatosensory (0% visual) attention; left: an AMI map of the responses overlaid on the MNI brain (blue denotes regions where alpha power decreased with increasing visual attention). The bar plot (right in panel A) shows the average, normalized, alpha power over subjects, where the power was measured from individual AMI minima in the right visual cortex over the four attention conditions. The blue line represents the linear fit of these data. **B**: Shows AAL region where significant linear modulation across conditions was observed. The region identified was the inferior occipital gyrus (marked in light blue, p=0.02, fdr-corrected). The modulation in this region is plotted in the bar graph (average normalised alpha responses across subjects) along with the line of best fit (blue line). **Panel C** shows the AMI map obtained when contrasting trials where subjects attended to only one modality (i.e. 100/0% condition) with those where attention was divided (i.e. 60/40% condition) overlaid on the MNI brain (red/yellow denotes regions where alpha power increases when attention is paid to a single modality compared with divided attention). The largest AMI effect to this contrast was in the right parietal area where an increase in alpha power is seen during 100%/0% attention conditions compared with 60%/40% conditions. The normalised alpha responses from the individual subjects’ peak locations are shown in the bar graph (panel C, right), revealing a quadratic (“u”-shaped) modulation. **D:** Shows AAL regions where significant quadratic modulation across conditions was observed. Both regions identified were in the parietal cortex (postcentral lobule (p-value = 0.003 (fdr-corrected)) and precuneus (p-value = 0.01 (fdr-corrected))). Interrogation of the alpha power responses in these regions, shown by the bar graphs (right panel of D), revealed a “u”-shaped across attention conditions in both regions. Error bars on all bar graphs denote the SEM across subjects for the normalised alpha responses.

To investigate whether differential alpha power modulation was observed in other brain regions in trials where attention was divided between modalities in comparison to those where attention was focused on one modality only, the AMI between the 100% (*cond 1* in Equation 1) and 60% (*cond 2* in Equation 1) attention conditions was computed. This contrast revealed a peak source location in the right parietal cortex, showing higher alpha power in the 100% than 60% attention conditions (Figure 3C, denoted by red colour). At this location, when all conditions were considered, a quadratic (“u-shaped”) alpha power response was observed (Figure 3C, right plot). AAL analysis confirmed this result, showing significant quadratic modulations of alpha power in two superior parietal regions: right post central lobule (p-value = 4×10^−5^, fdr-corrected) and right precuneus (p-value = 0.01, fdr-corrected). Visual inspection of the alpha power across conditions showed that significantly lower alpha power was induced in these regions when attention was divided between modalities than when subjects paid attention to only one modality (Figure 3D; left). No significant linear modulations were seen in these regions.

#### Experiment 2

In this second experiment AMI analysis in source space (Equation 1), identified a maximum in left and a minimum in right visual cortices when contrasting trials where subjects attended 100% to the left (*cond 1* in Equation 1) with trials where subjects attended 100% to the right (*cond 2* in Equation 1) side of the visual field (Figure 4A). Alpha power at the peak in the left visual cortex increased with increasing attention to the left visual hemifield, whereas the response in the right visual cortex showed a decrease in alpha power. For increasing attention to the right visual hemifield, the opposite was observed. Hence, a decrease in alpha power could be observed over visual areas contralateral to the focus of attention whereas an increase in alpha power was present over visual areas of the hemisphere ipsilateral to attention. These responses were combined by inverting the responses across conditions measured from right hemisphere and then averaging with those measured from left hemisphere. The results of this analysis are shown in the bar plot in Figure 4A, and suggest a linear modulation of alpha power by attention. The fitting analysis in AAL regions confirmed this observation, showing that with increasing attention, alpha power decreased linearly over visual areas of the hemisphere contralateral to the focus of attention: significant linear fits were found in the angular gyri (p= 0.03, fdr-corrected) and superior occipital gyri (p= 0.03, fdr-corrected), as shown in Figure 4B.

**Figure 4:**
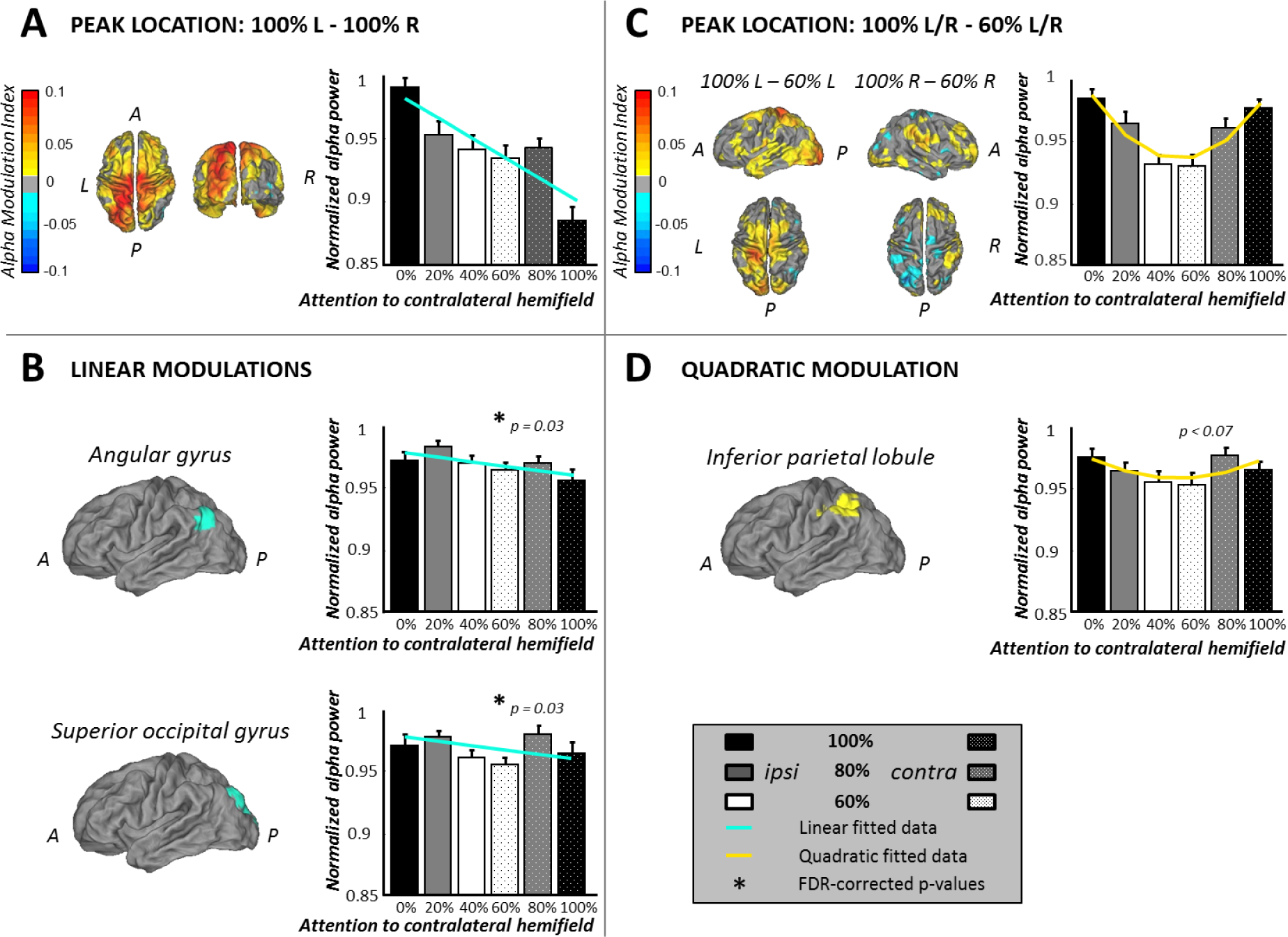
Source analysis results of Experiment 2. (**A**) shows the AMI map when contrasting conditions 100% attention left vs. 100% attention right (left side of the panel) revealing an increase (red/yellow colour) in alpha power over left visual and parietal areas for the 100% attention left condition compared with the 100% attention right condition (the contrast 100%R-100%L would just be the inverse of this AMI map). The bar plot shows the averaged, normalized, alpha power extracted from individual peak locations for the different attention conditions, showing decreasing alpha power values in the hemisphere contralateral to the location of increased spatial attention (data were extracted from maxima in the left and minima in the right visual cortex for attention 100% left and 100% right, respectively. Data extracted from the right visual peak location were inverted across conditions to be averaged with those of the left visual peak location). The light blue line shows the linear fit of the data. (**B**) shows the results of the AAL analysis revealing the angular gyrus (p= 0.03, FDR-corrected) and the superior occipital gyrus (p= 0.03, FDR-corrected) as the regions with a significant linear modulation of alpha power across the attention conditions (regions shown in light blue on the MNI brain). Bar plots show the alpha power over all conditions, again combined for the right and left hemisphere, the line of best fit is shown in light blue. (**C**) shows the AMI map when contrasting the attention conditions where participants attended to only one side of the visual field (100% L/R) with those when they divided their attention between left and right hemifields (60% L/R) overlaid on an MNI brain. The far left images show the responses to attention modulation to the left visual field, whilst the central brain maps show the same modulations with attention to the right visual field. The AMI maps show increase over ipsilateral parietal and visual areas to that side where attention is paid when attention is directed fully to that spatial location (100% condition) compared with divided between locations (60% condition). The bar plot (panel C) shows averaged normalized alpha power variation over attention conditions extracted from individuals’ peak locations in left and right parietal cortex, as applicable, with those responses from the right hemisphere inverted to be combined with left hemisphere responses. The yellow line depicts the quadratic fit to the data, outlining the seemingly “u”-shaped quadratic alpha power modulation. (**D**) shows the results of the AAL analysis with a trend of a quadratic modulation over the inferior parietal lobule (p= 0.07, FDR-corrected). The bar plot shows the alpha power over all conditions, again combined for the right and left hemisphere, the line of best fit is shown in yellow. Error bars on all bar graphs denote the SEM across subjects for the normalised alpha responses.

The AMI maps comparing the conditions 100% attention (*cond 1* in Equation 1) and 60% attention (*cond 2* in Equation 1) revealed maxima in the parietal cortex in the hemisphere ipsilateral to where visual attention was directed (Figure 4C, AMI maps), indicating higher alpha power in the 100% attention conditions than the 60% attention conditions. Interrogation of these responses over all conditions showed a quadratic (“u”-shaped) alpha power modulation pattern, as shown in the bar plot in Figure 4C. Further interrogation using the AAL analysis showed that a trend (p<0.07, FDR-corrected) for a quadratic fit was observed over the parietal region inferior parietal lobule (IPL). Visual inspection of the alpha power across conditions for this AAL region, revealed that the quadratic fit was a “u”-shape (Fig. 4D), as seen in the peak analysis (Fig. 4C) and similar to that seen for the multi-modal paradigm show in Figure 3C&D.

## DISCUSSION

Numerous EEG/MEG studies showed that posterior alpha power is modulated by attention. However, it is unclear whether these alpha power modulations reflect one or several attentional mechanisms. Here we show using EEG source analysis (LCMV beamformer) that in uni- and multimodal attention tasks, alpha power is differently modulated by attention in occipital and parietal areas (Figure 5).

**Figure 5:**
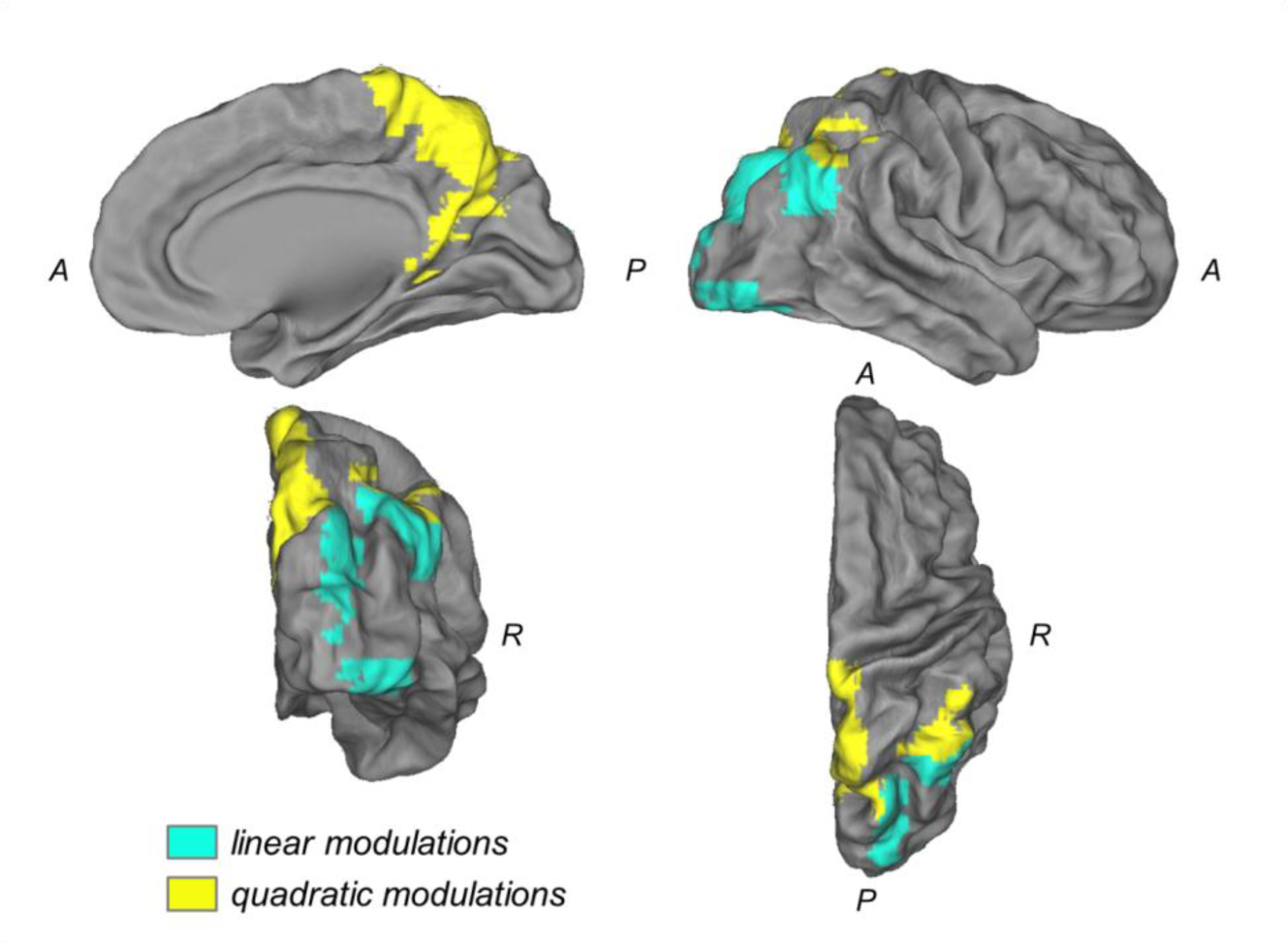
Summarizing Alpha power modulation effects with attention. To simplify, all effects are shown on the right hemisphere. AAL regions showing significant linear (light blue) and significant /trend quadratic (yellow) modulations of alpha power. A, P, and R stand for anterior, posterior, and right, respectively.

We found two alpha sources, visual and parietal, which can be separated spatially and experimentally. These sources are both modulated by attention, but play different functional roles depending on behavioural demands. The visual alpha source showed linear power decreases with increasing attention to visual stimuli or a given location, thus indicating the location of attention, i.e. the visual Spotlight of Attention (Posner et al., 1980; Crick, 1984; Eriksen and Yeh, 1985). In contrast, the parietal alpha source was quadratically modulated by attention showing lower alpha power when attention was divided, between modalities or spatial locations, rather than focused on either. Thus we suggest the parietal alpha source likely indicates attentional effort. Regions showing significant linear (light blue) and quadratic (yellow) alpha power modulations observed in Experiment 1 and 2 are summarized in Figure 5.

On the behavioural level, we have replicated previous findings (Gould et al., 2011) and show a robust effect of attention, modulating significantly accuracy and reaction times in both experiments where higher accuracy and lower reaction times are present when attention is focused on one modality (Experiment 1) or spatial location (Experiment 2).

### Functional significance

The two tasks used in this study show similar clustering of alpha activity in visual and parietal areas (Fig 5). This suggests that the two alpha sources are a general phenomenon of attention rather than specific to the experimental task. If our assumption is true, the imprecise region of interest that has previously been reported as *parieto-occipital* is actually composed of two distinct brain sources that act in different ways. We hypothesise that the sensory-specific source reflects “the visual spotlight of attention” and is controlled by top-down processes coming from a parietal alpha source which in turn is modulated by attentional effort towards the task.

Previous fMRI studies showed that both visual and parietal regions show an increased BOLD response in the hemisphere contralateral to the direction of visual spatial attention (Sylvester et al., 2007; Bressler et al., 2008; Lauritzen et al., 2009), reflecting increased cortical excitability. Using Granger Causality, Bressler et al. further showed that the FEF and IPS, both part of the dorsal attention network, were responsible for driving neural activity in early visual areas by top-down control (Bressler et al., 2008). Other studies obtained similar results (Ruff et al., 2008; Marshall et al., 2015; Popov et al., 2017). Since EEG alpha activity and the BOLD signal are widely reported to be negatively correlated (Goldman et al., 2002; Laufs et al., 2006; Scheeringa et al., 2011), an increase in BOLD signal over contralateral visual and parietal areas in a visual spatial attention task could be related to a decrease in alpha activity over the same regions, which would agree with the findings of this study. Capotosto et al. hypothesized that top-down control from frontal and parietal areas mediates the occipital alpha rhythm and therewith the level of inhibition (Capotosto et al., 2009). Using rTMS to inhibit the previously identified regions FEF and IPS (Bressler et al., 2008), Capotosto et al. showed increased reaction times and decreased accuracy for target detection. Furthermore, they demonstrated that this inhibition abolished the pre-stimulus alpha-desynchronization which can be typically observed over parietal and occipital electrodes contralateral to attention and concluded that this disruption in top-down control of the visual alpha rhythm led to a decrease in visual identification (Capotosto et al., 2009). However they were not able to identify what neuronal rhythms from IPS and FEF were causing this top down control of the occipito-parietal alpha rhythm from their experiment.

Previous studies support the idea for spatially distinct visual and parietal alpha sources mediating attention and visual perception. Van Dijk et al. (2008) showed that low pre-stimulus parietal alpha power was advantageous for visual discrimination. They concluded that this parietal alpha source regulates alpha power in low-level visual areas via top-down control. With our results we could re-interpret their findings and conclude that the parietal alpha power modulation rather reflects the attentional state than the level of inhibition of the occipital cortex. Thus, low parietal alpha power would indicate a state where the subject is engaged in the task, leading to the recruitment of top-down attention and an increase in detection performance. Another example is a recently published EEG study (Gulbinaite et al., 2017) on the triple-flash illusion, where a third visual flash is perceived upon presentation of only two stimuli. The illusion comes about when presenting the second stimulus after a specific interval; the authors could show that this interval and the illusory percept correlated with the individual alpha frequency at parietal but not occipital sources. In line with these findings, a recent intracranial EEG study shed more light onto these distinct alpha sources and their directionality, supporting the view of a top-down control of occipital alpha by parietal areas (Halgren et al., 2017). The authors recorded resting state data on epilepsy patients and found evidence for alpha generators in the parietal cortex. They further showed that alpha acts like a traveling wave, propagating in space from parietal to occipital brain regions (Halgren et al., 2017). Albeit the important evidence for the existence of two spatially distinct alpha sources, none of the above presented studies could experimentally dissociate them into occipital and parietal sources.

Crucially we extend these previous studies by showing that there are two distinct alpha sources which are modulated differentially by attention in two different sets of experiments and are thus likely to have different functional roles. This data adds to a growing body of evidence that there are multiple alpha sources present during a cognitive task with distinct roles (Nunez et al., 2001). Alpha oscillations have gained much interest in neuroscientific research and their image has changed from reflecting a passive idling state (Pfurtscheller et al., 1996) to actively regulating inhibition in the service of cognition (Klimesch et al., 2007; Palva and Palva, 2007; Jensen and Mazaheri, 2010; Mathewson et al., 2011). Given the ubiquity of alpha oscillations in the human brain, it makes sense to assume that the role played by alpha is a very general one like gating neural activity. If this assumption is true, then we should be able to dissociate different alpha oscillations in a complex cognitive task that recruits a number of cortical assemblies controlled by alpha. Our data represents such evidence where we spatially and experimentally dissociate an occipital/ventral parietal from a more superior parietal alpha source in two experiments requiring a complex interaction between top-down and bottom-up attention processes. Previous studies focused on the role of alpha in gating low level sensory information (Jensen et al., 2012). We add to this literature by showing that alpha not only indicates the locus of attention, but also the recruitment of higher order areas, which arguably control the shift of attention to lower-order, primary sensory locations/modalities. Our results open up the avenue for future non-invasive human EEG studies to investigate how alpha oscillations in these two regions coordinate their activity to implement attentional shifts, which so far has mostly been addressed by invasive animal recordings (von Stein et al., 2000; Buffalo et al., 2011; van Kerkoerle et al., 2014).

EEG source localization relies on whether the assumptions of its algorithm are met by the data. Our EEG results are corroborated by a separate, high resolution fMRI study conducted by our group at ultra-high field (7T) on a sub-sample of the same subjects (7/10 participants also performed Experiment 1), using the same multi-modal task as in Experiment 1. This study (Aquino et al., 2018) also reveals two fMRI sources modulated by attention: i) quadratic BOLD-response modulations over parietal areas when contrasting attentional effort (100 vs. 60%) and ii) linear modulations over visual areas when contrasting the location of attention (100 vs. 100%). Due to the superior spatial resolution of fMRI, these results strongly suggest that the two alpha band sources measured with EEG are indeed distinct sources which need to be considered separately.

### Conclusions

We show that two spatially distinct alpha sources execute different roles in uni- and multi-modal attention: i) a parietal source, modulated by attentional effort showed significantly lower alpha power when subjects divided their attention which potentially exerts top-down control on alpha oscillations over lower-level visual areas, ii) a visual alpha source that reflects the current spotlight of visual attention showing a significant linear power decrease with increasing attention to visual stimuli, possibly driven by top-down control from parietal alpha sources. Given that such a top-down control has been shown to exist (i.e. Bressler et al., 2008; Ruff et al., 2008; Capotosto et al., 2009), we hypothesize a similar top-down regulation from parietal towards visual areas, however, further exploration is needed to confirm this hypothesis.

## Acknowledgements

This research was funded by a Leverhulme Trust Research Project Grant to KJM.

